# An expanded urine culturing workflow to cultivate and characterize diverse urobiome isolates

**DOI:** 10.64898/2026.08.24.746228

**Authors:** Freya Darling Eriksen, Marleen D. Hekker, Corné van der Zeeuw, Tyrsa Veld, Gerald Wittenaar, Maria Jové Casals, Kaity-Lynn Buiting, Jolanda K. Brons, Pablo Gallardo Molina, Michael F. Seidl, Rampal S. Etienne, Thomas Hackl, Alan J. Wolfe, Janneke H.H.M. van de Wijgert, Marjon G.J. de Vos

## Abstract

**Background:** Despite increased recognition of the urinary tract’s diverse resident microbiome (i.e., the urobiome) in postmenopausal women, the roles and functions of these microbes remain largely unknown. Further empirical research is needed to understand the physiology, interactions, and antibiotic resistance evolution of urobiome members with pathogenic potential. However, experimental work relies on viable, culturable isolates. Standard urine culturing practices are designed for identifying a narrow set of known urinary microbes, and are thus poorly suited for cultivating taxa from the resident urobiome. Here we expand the urine culturing toolkit to reliably recover diverse urobiome taxa for downstream empirical research.

**Methods:** Urine samples collected from postmenopausal women with recurrent urinary tract infections were shipped at ambient temperature to a central point for culturing. Microbial viability was maintained using boric acid preservative tubes during multi-day transport of sample aliquots. Selective media incubated under specialized conditions were used to promote recovery of diverse urobiome members, including fastidious taxa. Under 5% CO_2_-enriched atmospheric conditions and with longer incubation times, we leveraged a chromogenic agar (UTIC) to further differentiate isolates based on colony color and morphology. We evaluated the workflow for its ability to isolate and characterize urobiome taxa, as determined by morphological differentiation and taxonomic identification.

**Results:** Across 108 urine samples, 6.3 ± 3.2 distinct isolates were recovered, with no detectable relationship between sample shipment duration and isolate richness. On chromogenic agar, colony growth and color intensity was improved with CO_2_-enriched atmospheric conditions and extended incubation times. We identified diverse taxa that are typically underrepresented in standard diagnostic culture and provide novel morphological characterizations for members of the genera *Actinotignum, Aerococcus*, *Facklamia*, *Lactobacillus*, *Latilactobacillus*, *Limosilactobacillus,* and *Streptococcus* species, which have not been previously described on UTIC chromogenic agar.

**Discussion:** Using this novel workflow, we recovered a diverse collection of urobiome isolates from urine samples shipped over multiple days. We also demonstrated the utility of a chromogenic agar for the visual differentiation of key urobiome taxa. While sequencing approaches have enhanced our understanding of urobiome composition, culturing is needed to investigate microbial interactions, virulence mechanisms, and antimicrobial susceptibility. This protocol adds to the growing toolkit for the cultivation of diverse urobiome isolates needed to support downstream empirical studies and advance urinary tract infection research.

## Background

Urinary tract infections (UTIs) are among the most common bacterial infections worldwide and place a disproportionate burden on women. One in two women will experience at least one UTI in their lifetime, and risk of developing recurrent UTIs increases significantly in postmenopausal women [1, 2]. Diagnosis and treatment of UTIs remains challenging due to nonspecific symptoms, culture-based detection limitations, and antimicrobial resistance [3–5].

Advances in sequencing techniques and enhanced cultivation methods such as Expanded Quantitative Urine Culturing (EQUC) have revealed a wide diversity of bacteria residing in the human urinary tract, now known as the urobiome [6–12]. Due to the longstanding misconception that urine is sterile in the absence of infection, UTI research and clinical diagnostics often focus on a limited set of species thought to be uropathogenic, while other urobiome members go undetected or are dismissed as contamination [1, 10, 13–15]

Recent work suggests an important role of the urobiome in UTIs. Many taxa previously overlooked by standard urine culturing have been linked to urinary tract symptoms [14, 16] and inflammatory biomarkers in urine [12], while network-based analyses have shown that clusters of co-occurring urobiome taxa may reflect relevant interactions underlying UTIs [1]. Importantly, ecological interactions between urobiome members have been shown to alter the growth, virulence, and antibiotic resistance of bacteria with uropathogenic potential [13, 15, 17–21].

Standard urine culturing methods target bacteria from a short list of recognized uropathogens especially *Escherichia coli* and including other Gram-negative bacteria [5]. However, modern DNA-based identification approaches and selective culturing methods have revealed a much greater diversity of bacteria than previously expected in urine of individuals with and without UTI [12, 22]. Notably, the presence of *E. coli* and other Gram-negative bacteria, even at high abundances, is not always predictive of UTI, as these bacteria can be present in both individuals who do and do not meet clinical UTI diagnostic criteria [1, 16, 22]. Thus, the current uropathogen-centric paradigm overlooks the urobiome, limiting our understanding of UTIs, and highlighting the need for further urobiome investigation.

Empirical UTI research is constrained by the available isolates obtained through standard culturing methods, leaving a wide diversity of urobiome taxa unexplored [23, 24]. Without cultured isolates to experimentally manipulate, their capabilities and ecological roles cannot be investigated. Future investigations of microbial physiology, interactions, and antibiotic resistance evolution will require new tools to culture microbes from the urobiome and manipulate those isolates *in vitro*. While sequencing technologies will remain a critical tool in urobiome research, our knowledge of “who is there” will become significantly more impactful when “what they do” is also known. Building and testing a more diverse urobiome isolate collection is thus an essential prerequisite to advancing the urobiome research field.

Over the past decade, enhanced cultivation methods such as EQUC in larger cohorts have helped to identify and isolate many urobiome taxa typically overlooked by standard methods [9, 25]. Here we present the development and application of a culturing workflow to capture diverse bacterial isolates from urine samples for downstream empirical studies. The aim of this workflow is to provide a practical tool to facilitate collaborative projects and geographically expand urobiome research.

In particular, we demonstrate the utility of Urinary Tract Infections Chromogenic agar (UTIC; Condalab) for the visual differentiation of taxa not previously characterized on UTIC. These include taxa considered to be emerging uropathogens [14], including members of the genera *Actinotignum, Aerococcus*, *Facklamia,* and *Winkia*, as well as taxa considered to be beneficial, including diverse lactobacilli. We also show that even during multi-day shipment durations at ambient temperature, not only known urinary pathobionts [26], but also diverse urobiome isolates can be captured from urine samples preserved in boric acid.

## Materials and Methods

### Urine Sample Collection and Study Cohort

Urine samples were collected as part of an interdisciplinary project (the UTI Revisited or UTIr project) aiming to identify microbial eco-evolutionary drivers of recurrent UTIs (RUTIs). Postmenopausal women suffering from RUTIs (defined as at least 3 symptomatic UTIs in the past 12 months) and postmenopausal controls without a RUTI history and no UTI in the past 12 months were recruited by the University Medical Center Utrecht, the Netherlands. Cohort recruitment, and the one-year extensive sample collection procedures have been previously described [27]. In brief, midstream voided urine samples were collected both at pre-scheduled time points in the absence of a UTI, and during UTI episodes. The start and end of UTI episodes were determined based on self-reported symptoms.

### Urine Sample Storage and Shipment

To preserve microbial viability and prevent overgrowth during transport, 4 mL of the urine sample was aliquotted immediately after collection by a study team member into a BD Vacutainer Plus C&S Urine Tube containing boric acid sodium borate/formate as preservative (Becton Dickinson BV, Drachten, Netherlands). Aliquotted samples were shipped the same day via post to the culturing site at the University of Groningen, the Netherlands, in medical post envelopes. Upon arrival, samples were immediately refrigerated at 4°C and the culturing protocol began within 24 hours.

### Workflow Overview

We established a workflow to process and culture urine samples shipped over multiple days. Boric acid preservative tubes allowed for samples to remain suitable for culturing, allowing for downstream processing of samples at the central culturing location. Upon arrival, samples were refrigerated at 4°C and processed within 24 h using the protocol described below. The workflow contained four stages: (i) plating aliquots of urine onto multiple selective agars at two volumes, (ii) incubating plates under agar-specific atmospheric conditions for 3-4 days, (iii) selecting and isolating morphologically distinct colonies onto a chromogenic agar for further differentiation, and (iv) generating pure cultures for long-term storage and taxonomic identification. This workflow enabled the culturing of diverse, morphologically distinct urobiome isolates from shipped urine samples.

### Media and Agar

Different types of nutrient agars were used to improve visualization of distinct colonies and support the growth of fastidious organisms. Four selective media were used: Columbia Nalidixic Acid agar (CNA; BD), MacConkey agar (MAC; Sigma Aldrich), De Man-Rogosa-Sharpe agar (MRS; Sigma Aldrich), *Gardnerella vaginalis* selective agar (GARD; Thermo Scientific), along with a non-selective Tryptic Soy agar supplemented with 5% sheep’s blood (TSA; Thermo Scientific). The CNA, MAC, MRS, and GARD media are selective for Gram-positive, Gram-negative, lactobacilli, and *Gardnerella,* respectively. CNA, MRS, and MAC agars were prepared from powder. MRS agar was prepared according to manufacturer instructions with the addition of Tween® 80 (MP; Biomedicals) and GARD agars were pre-poured by the manufacturers. Agar plates were stored at 4°C for up to 4 weeks prior to usage in workflow or until the manufacturer-defined expiration date. Following the first incubation round, morphologically distinct colonies from each nutrient agar were selected and streaked onto CondaChrome® UTIC agar (UTIC; Condalab). Colonies were then inoculated into Tryptic Soy broth (TSB; Sigma Aldrich) to make liquid cultures for subsequent isolate storage.

### Urine Culturing

Urine samples were directly plated at two volumes (2µL and 100µL) onto CNA, MAC, and MRS nutrient agars. TSA and GARD agars were inoculated with only the lower volume of urine (2µL), due to frequent overgrowth on the relatively nutrient-rich agars. Sterile glass 3 mm beads were used to spread the appropriate volume of urine across each agar plate. CNA plates were incubated at 37°C in 5% CO_2_ and 18% O_2_; MAC agar plates were incubated aerobically at 37°C; MRS, TSA, and GARD agar were incubated at 35°C with one Oxoid AnaeroGen 2.5L sachet (Thermo Scientific) in an anaerobic box to generate a low-oxygen atmosphere (CO_2_ between 8% and 14%, O_2_ < 1%). All selective agars were incubated in their respective conditions for 3-4 days. On each selective agar plate, colony forming units were estimated at 5 different levels of density (0, 1-10, 10-50, 100+, TD) ranging from no visible growth (0) to too dense (TD) to count. Morphologically distinct colonies from each nutrient agar were selected and subsequently streaked onto UTIC agar and incubated at 37°C in an enriched CO_2_ atmosphere (5% CO_2_, 18% O_2_) for 3-4 days.

### Isolate Selection and Identification

Colony morphologies were assessed after full pigment development on UTIC agar (37°C in 5% CO_2_; 2-4 days incubation). Descriptions reflected predominant and consistent visual characteristics including color, size, opacity, and edge morphology. Color categories included main color values, as well as two shade possibilities (light or dark) in addition to each main color. Size categories were defined as punctiform (<1 mm), small (1–2 mm), medium (2–4 mm), or large (>4 mm). Isolate numbers were estimated based on the number of morphologically distinct isolates observed on UTIC agar. Additionally, UTIC agar served as a contamination check to ensure that colonies from the nutrient agars were pure for subsequent storage and taxonomic identification. Based on their visual characteristics on UTIC agar, all morphologically distinct colonies were inoculated into 8mL TSB and incubated at 37°C in 5% CO_2_ for 2-4 days. Colonies from the first incubation round that did not grow on UTIC agar were inoculated directly from their origin agar into TSB in the next culturing step. From the liquid cultures, 750µL of each isolate was stored in an equal volume of 50% glycerol, with a final concentration of 25% glycerol (v/v) (Sigma Aldrich) in 2mL cryo tubes at -80°C. Taxonomic identification of isolates was determined using either matrix-assisted laser desorption ionization time-of-flight mass spectrometer (MALDI-TOF MS) or 16S rRNA gene Sanger sequencing. All isolates, regardless of sample shipment duration, were included for subsequent isolate selection, identification, and morphological descriptions. All morphological observations were based on isolates collected in the present study, with the exception of one *Lactobacillus crispatus* isolate obtained from an asymptomatic woman in a previous study [28]. This additional isolate was included because *L. crispatus* was confirmed only once in our samples, thus insufficient isolates were available for morphological description.

### Identification of isolates by MALDI-TOF MS

Individual colonies isolated from UTIC agar were deposited onto the MALDI-TOF target plate using a sterile toothpick. Each spot was coated with 1 µL of HCCA matrix (Bruker Daltonics, Germany) and processed by matrix-assisted laser desorption/ionization time-of-flight mass spectrometry (MALDI-TOF MS; MALDI Biotyper®, Bruker Daltonics, Germany; library version 5.1.420) for bacterial identification. Each isolate was biotyped in duplicate.

### Identification of isolates by 16S rRNA gene Sanger sequencing

Isolates grown on UTIC agar were subjected to colony PCR for 16S rRNA gene amplification. A single colony was suspended in 50 µL sterile Milli-Q water; 3 µL of the undiluted suspension and a 10-fold diluted suspension were used as PCR templates. Nearly full-length 16S rRNA genes were amplified using the universal primers B8F (5′-AGAGTTTGATCMTGGCTCAG-3′) and 1492R (5′-GGTTACCTTGTTACGACTT-3′) [29].

The PCR mixtures were composed as follows: 2.5 μL of 10x PCR reaction buffer (Roche, Basel, Switzerland) supplemented with an additional 0.8 mM MgCl₂ (Merck, Darmstadt, Germany), 2% dimethylsulfoxide (DMSO, Merck, Darmstadt, Germany), 20 µg/mL bovine serum albumin (BSA, Merck, Darmstadt, Germany), 200 µM of each dNTP (Merck, Darmstadt, Germany), 0.2 µM of each primer (Eurogentec, Seraing, Belgium), and 1 U of Taq DNA polymerase (Roche, Basel, Switzerland). The final volume was adjusted to 25 μL with molecular biology grade water (Thermo Fisher Scientific, Waltham, USA) in a 0.2 mL microcentrifuge tube. Amplification was performed in a Mastercycler Nexus thermal cycler (Eppendorf, Hamburg, Germany), programmed as follows: initial denaturation at 95 °C for 10 min; 35 cycles of 95 °C for 1 min, 51.8 °C for 30 s, and 72 °C for 2 min; followed by a final extension at 72 °C for 5 min. PCR products were analyzed via 1.0% (w/v) agarose gel electrophoresis, stained with ethidium bromide (1.2 mg/L), and visualized under ultraviolet light [30].

Amplicons were submitted to BaseClear (Leiden, the Netherlands) for 16S rRNA gene Sanger sequencing [31]. Raw reads were trimmed using the *sangeranalyseR* R package, [32] and trimming quality was assessed by manual inspection of chromatograms. Sequences were classified using two complementary approaches. First, sequences were aligned and classified using the SINA aligner (v1.2.12) through the SILVA web interface [33] querying SILVA, RDP and LTP databases. Second, ambiguous nucleotides were replaced with “N” using the *Biostrings* R package [34] after which sequences were classified using the BLASTn tool (v2.17+) against the NCBI nucleotide database. For sequences unable to be taxonomically assigned by SINA/SILVA, the BLASTn-derived classification was used. In all other cases, taxonomic assignments were concordant with both approaches.

### Data Analysis

Shipment duration (defined as the number of days between urine sample collection and initiation of processing) was treated as a categorical predictor. Overall isolate counts were summarized using the mean ± standard deviation (SD) and median. The majority of samples were processed after shipment durations of 2-7 days, and were included in the isolate richness data analysis; samples with longer shipment durations were excluded. Due to the count nature of the data and overdispersion, the association between shipment duration and isolate richness was further evaluated using a negative binomial regression. Shipment duration was modeled as a categorical variable with the 2-day group specified as the reference category, representing the shortest possible shipment duration. The overall significance of shipment duration was assessed using likelihood-ratio chi-square tests. Statistical analyses were performed with R version 4.4.3 (R Core Team, 2025). All isolates, regardless of shipment duration, originating from CNA (*n* = 403), MRS (*n* = 100), and MAC (*n* = 61) agars were included in analysis of colony color and morphological characterizations.

## Results

### Obtained Isolates

We successfully recovered a diverse set of urobiome isolates using the workflow described above. The recovered isolates included both Gram-positive and Gram-negative taxa commonly associated with the urogenital microbiome [5, 24, 35, 36] including organisms of established clinical relevance as well as taxa that remain underrepresented in standard diagnostic culture including *Actinotignum*, *Aerococcus*, *Facklamia*, and *Winkia* species, as well as various lactobacilli (Table 1).

**Table 1.** Morphological descriptions of Gram-positive urobiome taxa not previously described on chromogenic (UTIC) agar, following incubation for 3-4 days at 37°C under 5% CO_2_ conditions. Taxa which have not yet been morphologically described may have similar appearances to those listed here. Taxa with genus-level morphological similarity are grouped in the same row, with the identified species contributing to the description listed below the shared genus.

| Taxa | Colony Color | Colony Morphology | Agar Origin |
| --- | --- | --- | --- |
| *Actinotignum* spp.<br>*A. schaalii*<br>*A. timonense* | Turquoise (Light) | Punctiform, scanty, slight central pigment intensification | CNA |
| *Aerococcus* spp.<br>*A. sanguinicola*<br>*A. urinae* | Cream | Punctiform-small, occasional irregular (non-circular) shape, central pigment intensification | CNA, TSA |
| *Enterococcus faecalis* | Turquoise (Light) | Medium, circular, diffused color on edges | CNA, MRS, TSA |
| *Enterococcus faecium* | Turquoise (Dark) | Medium, circular, diffused color on edges | CNA, MRS |
| *Facklamia hominis* | Magenta | Small, circular, diffused color on edges | CNA, TSA |
| *Lactobacillus crispatus* | Turquoise (Dark) or Magenta | Punctiform-small, circular, solid color throughout, two observed color morphs | TSA |
| *Lactobacillus gasseri* | Turquoise (Light) | Small-medium, lobate, central pigment intensification | CNA, MRS, TSA |
| *Lactobacillus jensenii* | Turquoise (Light) | Punctiform-small, lobate, slight central pigment intensification | CNA, MRS, TSA |
| *Latilactobacillus sakei* | Magenta | Punctiform-small, circular, serrated edges, central pigment intensification, | CNA |
| *Limosilactobacillus reuteri* | Magenta | Small, circular-irregular, central pigment intensification | MRS |
| *Streptococcus* spp.<br>*S. anginosus*<br>*S. vaginalis* | Turquoise (Dark) | Small-medium, circular, diffused color on edges | CNA |

### Growth on Chromogenic Agar

Unique colony morphologies of urobiome isolates grown on UTIC agar (37°C; enriched CO_2_ atmosphere) were used to differentiate taxa for both isolate collection and downstream empirical applications. Using these methods, we show that reproducible colony colors and morphologies can be obtained on UTIC agar for diverse urobiome isolates, including fastidious taxa previously undescribed on this chromogenic agar (Figure 1). We report the correspondence between newly-described colony morphologies on UTIC agar and representative taxa, excluding previously-established color associations (Table 1; Table S1).

**Figure 1.**
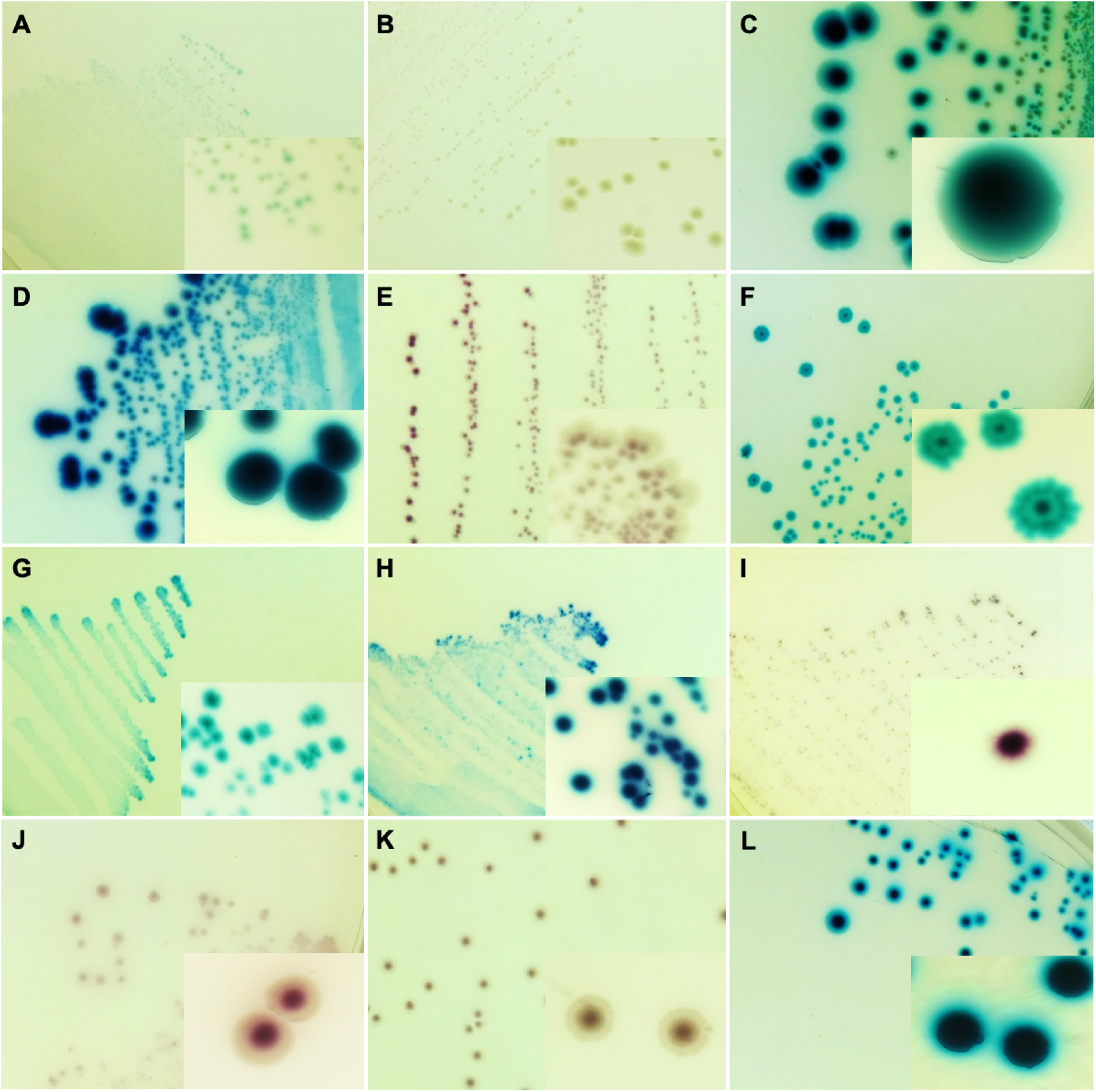
Morphologies of select isolates grown on chromogenic agar (UTIC), following incubation for 3-4 days at 37°C under 5% CO_2_ conditions. Isolates pictured are identifiable at the genus or species level. Additional morphological characteristics and descriptions can be found in Table 1. Each image shows the colonies as viewed with a naked eye and a macro photograph in the bottom right corner. **A** *Actinotignum* spp. **B** *Aerococcus* spp. **C** *Enterococcus faecalis*. **D** *Enterococcus faecium.* **E** *Facklamia hominis.* **F** *Lactobacillus gasseri / paragasseri.* **G** *Lactobacillus jensenii.* **H** *Lactobacillus crispatus* (dark turquoise morph). **I** *Lactobacillus crispatus* (magenta morph). **J** Limosilactobacillus reuteri. **K** *Latilactobacillus sakei.* **L** *Streptococcus anginosus / vaginalis*

The use of CO_2_-enriched atmospheric conditions and longer incubation times for isolates on UTIC agar produced noticeably more vibrant and robust colony colors compared to those incubated under standard conditions. For instance, the genus *Lactobacillus* had been previously described on UTIC agar as scanty and light blue [37], which differs from our observation of robust growth and vibrant colors when grown under more suitable conditions. We observed that the standard 18-24 hours of incubation time was not sufficient for many isolates to develop a color on UTIC agar and that many fastidious urobiome isolates failed to grow on UTIC agar in standard aerobic conditions.

UTIC agar produced distinct and reproducible colony morphologies across a broad range of taxa. Identifications of select taxa could be made using morphology alone (Table 1; Figure 1). Morphologies tended to be similar at the genus level, with the exception of species from the genera *Corynebacterium* and *Staphylococcus* which showed higher morphological diversity (Table S1). The phylogenetic conservation of colony morphology was also observed for many lactobacilli, and corresponded with the group’s recent re-classification into multiple distinct genera including *Lactobacillus*, *Latilactobacillus, Limosilactobacillus*. An exception to the conservation of morphology at the species level applied to *Lactobacillus crispatus*, for which two distinct color morphs were observed (Figure 1H and 1I).

Across all samples, 671 morphologically distinct urobiome isolates were obtained. The proportion of distinct color morphologies of isolates obtained from CNA (*n* = 403), MRS (*n* = 100), and MAC (*n* = 61) differed depending on the selective agar origin plate (Figure 2). Isolates from MAC agar yielded mainly expected chromogenic appearances of previously-described Gram-negative bacteria when subsequently grown on UTIC agar, including the red colony morphology characteristic of *E. coli* (*n* = 7) Additional isolates originating from MAC agar exhibited appearances on UTIC agar consistent with other Enterobacterales, including *Klebsiella* (dark blue; *n* = 2) and *Proteus* (yellow to orange; *n = 3*) spp., although at a lower proportion.

**Figure 2.**
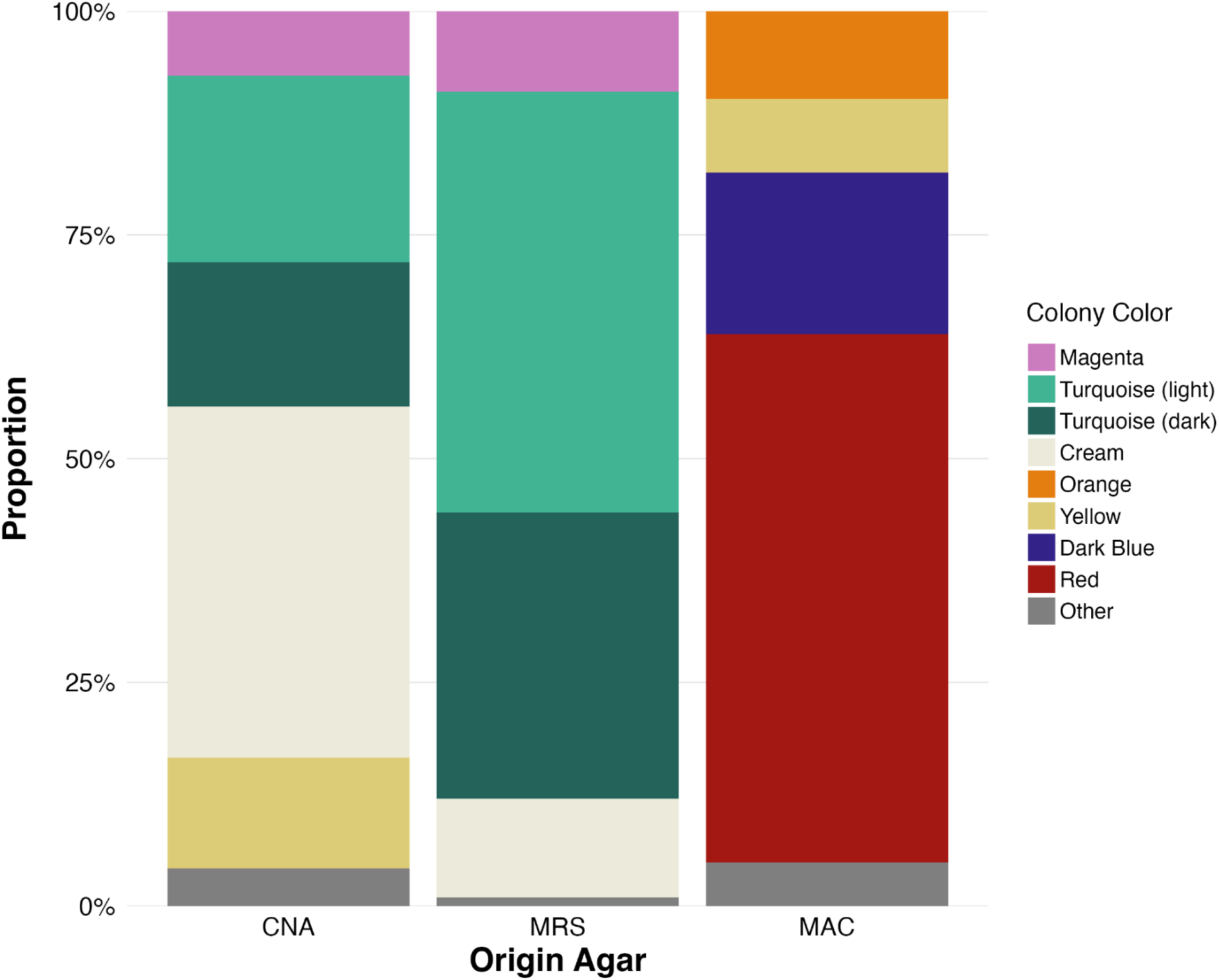
Proportion of distinct colony color morphologies observed on UTIC agar following transfer from three selective origin agars (MAC, CNA, and MRS). Color morphologies are recorded per isolate. CNA-derived isolates (*n* = 403) presented the greatest overall morphological diversity and were predominantly turquoise and cream. MRS-derived isolates (*n* = 100) were predominantly turquoise with the greatest proportions of magenta morphologies. MAC-derived isolates (*n* = 61) were predominantly red, characteristic of *E. coli*. ‘Other’ denotes colors comprising less than 0.25% of all total isolates in each selective agar of origin. Not all isolates characterized by color have been taxonomically identified by MALDI-TOF or 16S sequencing. Figure colors are adjusted to be color-blind-friendly and do not replicate exact color shade or hue as seen on UTIC agar.

Isolates originating from CNA agar, which is selective for Gram-positive bacteria, exhibited the highest isolate richness among samples, and greatest diversity as defined by the number of morphologically distinct isolates on UTIC agar. CNA-derived colony colors were mainly cream, corresponding to staphylococci, among others.

Isolates from *Gardnerella*-selective agar that subsequently grew on UTIC agar were assumed to not be *Gardnerella*, due to this taxon’s known inability to grow on UTIC agar. Isolates originating from *Gardnerella*-selective agar were therefore excluded from morphology characterizations on UTIC agar. The single *L. crispatus* isolate obtained in this study was derived from TSA agar. Distinctive morphological characteristics of taxa confirmed by taxonomic identification with MALDI-TOF or 16S rRNA gene sequencing are described in Table 1.

Both CNA-derived and MRS-derived isolates included a high proportion of turquoise colonies, corresponding to enterococci, streptococci, and other Gram-positive taxa. In the present study, we observed Streptococcus spp. (*n* = 15) displaying significantly more vibrant turquoise colors compared to previous descriptions of this taxon. We also found that *E. faecium* and *E. faecalis* can be clearly and consistently differentiated by colony color, with *E. faecalis* (*n* = 15) producing distinctly lighter turquoise colony colors compared to the darker turquoise of *E. faecium* (*n* = 4) (Figure S1).

Less common and previously undescribed morphologies were observed, including the magenta *Facklamia hominis* (*n* = 3) and light turquoise *Lactobacillus jensenii* (*n* = 8). Beyond colony color, other morphological characteristics also allowed for differentiation of taxa. For instance, *Aerococcus* species (*n* = 9), *Actinotignum* species (*n* = 2), *Latilactobacillus sakei* (*n* = 1), and *Limosilactobacillus reuteri* (*n* = 2) could be identified by their characteristic central pigment intensification, while *Lactobacillus gasseri* (*n* = 13) isolates could be differentiable by their distinct lobate edges (Table 1 and Figure 1).

### Preservation of Urobiome Isolates During Shipment

The mean number of days from shipment to start of processing was 4 ± 1.7, with a median of 4. We successfully cultured a broad range of urobiome isolates from urine samples preserved in boric acid and shipped at ambient temperature for multiple days. Across all urine samples included in analysis (*n* = 108 samples), the mean number of distinct isolates obtained from the final culturing step on UTIC agar was 6.3 ± 3.2, with a median of 6 (range 1-16 isolates). Isolate richness was not affected by shipment duration and did not differ for any of the shipment durations up to and including 7 days, with the exception of 5 days, which had significantly lower isolate richness compared to 2 days (*p* < 0.05) (Figure 3). Even when all shipment durations were included, the median isolate number remained 6 (Figure S2). Visible overgrowth of bacteria in samples was not observed.

**Figure 3.**
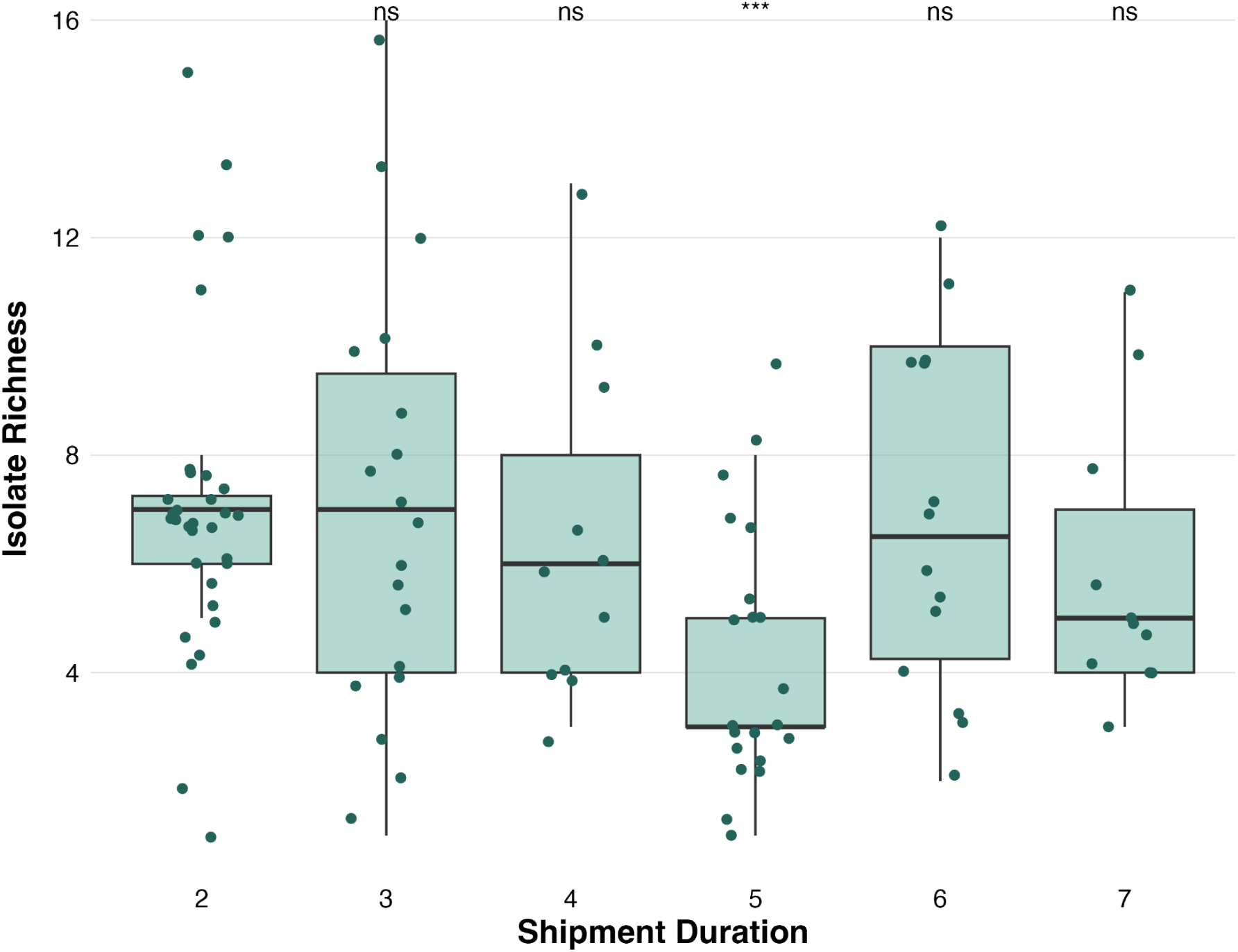
Isolate richness at different shipment durations across urine samples. Excluding samples with shipment durations longer than 7 days. Isolate richness is defined as the number of morphologically distinct isolates as described on UTIC agar obtained from one urine sample (*n* = 108 urine samples). Shipment duration is defined as the number of days between urine sample collection and starting the culturing workflow. (ns = not significant, ***p < 0.001).

## Discussion

Empirical research into the urobiome requires the recovery of viable bacterial isolates for downstream experimental work, including functional characterizations, comparative growth assays, and antimicrobial susceptibility testing. Standard culturing tools often fail to grow fastidious organisms, overlooking many key urobiome taxa.

Diagnostic urine culturing practices often require fresh urine, presenting a logistical constraint for research applications, particularly when sample collection and laboratory processing need to occur at different locations or when extended workflows are required. One remedy to this is the use of preservatives, such as boric acid. Boric acid has routinely demonstrated the recovery of urinary pathobionts from urine samples during short transit times (24-48 hours) [10, 26, 38–40]. A systematic evaluation of boric acid preservation in urine samples is still needed, both for its utility in the recovery of diverse urobiome isolates, particularly Gram-positive taxa, and for the preservation of urine longer than 24 hours [41].

More recently, boric acid has been increasingly applied to urobiome research, preserving urine samples prior to the development of EQUC and other downstream analyses [12, 42, 43]. Here, we utilized boric acid tubes to preserve urine samples during multi-day transport for subsequent culturing. In addition to previous work [12, 42–44], our findings support the utility of boric acid for the preservation and cultivation of diverse urobiome isolates, in addition to uropathogens. These results indicate that boric acid preservation supported the recovery of viable urobiome isolates during multi-day ambient-temperature shipment, with no apparent relationship between recovered isolate richness and shipment duration. However, further taxonomic comparisons between cultured isolates and taxa that are present in the urine sample but did not grow in culture will be needed to determine if specific urobiome taxa are lost during longer transit durations or prolonged exposure to boric acid.

The enhanced culturing method presented here requires multiple days of processing, different incubation steps, and multiple growth conditions, which disqualifies its use as a diagnostic tool when prompt results are needed. However, the preservation of diverse urobiome isolates in urine samples without the need for freezer storage could have many research applications. Longer-term sample preservation would significantly expand the geographic scope of urobiome research, allowing for more collaborative research opportunities. The collection and shipment of urine samples from various locations could also pave the way for future citizen science projects, allowing for the inclusion of more demographically diverse cohorts.

Chromogenic CondaChrome® UTIC agar (Condalab) was originally developed for the identification of microbes considered to be uropathogens, namely *E. coli*, *Klebsiella* spp., *Pseudomonas aeruginosa, Proteus mirabilis, Salmonella typhimurium Staphylococcus aureus*, and *E. faecalis* [45]. More recently, an updated CondaChrome® Pro UTIC agar was developed for the additional detection of *Citrobacter freundii, Streptococcus agalacticae,* and *Staphylococcus saprophyticus*. The application of UTIC agar on the broader urobiome, along with the implementation of specialized culturing conditions, has not been previously described. Here, we demonstrate the utility of UTIC agar (original media formulation) for the differentiation of diverse and fastidious urobiome taxa, including members of the genera *Actinotignum, Aerococcus, Facklamia*, *Lactobacillus*, *Latilactobacillus*, *Limosilactobacillus, Streptococcus,* and *Winkia*.

We observe that UTIC agar is able to reliably differentiate between different genera, even those only recently defined using genomic tools, as in the case of lactobacilli. Notably, *Limosilactobacillus reuteri,* which was only very recently split from its original *Lactobacillus* genus [46], yields a completely different color (magenta) on UTIC agar compared to the taxa from its former genera *Lactobacillus gasseri* and *Lactobacillus jensenii* (turquoise). The one observed exception to this genus-level morphology conservation was *Lactobacillus crispatus*, which presented both dark turquoise and magenta color morphs (Table 1; Figure 1). Only one *L. crispatus* isolate was cultured in the present study. Therefore, an additional *L. crispatus* isolate collected in a previous study [28] was included for the purpose of describing this taxon’s morphology. Further work leveraging expanded culturing methods are needed to gain further insight into the colony morphologies of the diverse lactobacilli, and other bacterial isolates cultured.

It has been observed using sequencing methods that *L. crispatus* is more abundant in urine and vaginal samples from premenopausal women compared to postmenopausal women [47]. The present study cultured urine from postmenopausal women, which could also have resulted in the lower number of verified *L. crispatus* isolates compared to the other identified lactobacilli. Additionally, the culturing and preservation methods may have not been ideal for this species, which may have led to markedly lower detection via culturing (Materials and Methods). We also demonstrate the ability to visually differentiate between isolates within the same genus on UTIC agar. Previously, *Enterococcus* spp. had been characterized at the genus level on UTIC agar. Here, we observe that *E. faecium* and *E. faecalis* can be clearly and consistently differentiated, with *E. faecium* producing distinctly darker colony colors compared to the lighter turquoise of *E. faecalis* (Figure 1; Figure S1). Future applications of UTIC agar for the differentiation of cultured lactobacilli, enterococci, and other diverse urobiome taxa should be investigated.

Differentiation of taxa based on morphology on UTIC agar as part of an EQUC-type protocol could, after rigorous further testing, potentially provide diagnostic value by identifying relevant urobiome taxa, particularly in settings where other means of identification (for example, with MALDI-TOF or sequencing) may not be feasible. UTIC agar could also serve as a tool for the visualization of urobiome taxa in various experimental applications, including co-growth interaction studies or synthetic urobiome communities.

Although sequencing-based methods offer powerful insights into urobiome composition, compositional data alone are insufficient without a mechanistic understanding of underlying urobiome dynamics. Investigations of virulence mechanisms, microbial interactions, host-effects, and antibiotic resistance all depend on the availability of viable, isolated, and culturable microbes. Additionally, the cultivation of taxa following multi-day sample shipment can geographically expand research efforts, providing opportunities for the expansion of study cohorts and more collaborative urobiome research projects.

## Conclusions

The workflow presented here enables the cultivation and characterization of diverse urobiome taxa, contributing to the culturing toolkit needed to support downstream empirical work into microbial interactions, antibiotic resistance, and the molecular mechanisms underlying virulence traits. We anticipate that investment in both sequencing and experimental approaches, along with the development of robust culturing tools, will enable continued progress in urobiome research.

## Abbreviations

CNA: Columbia Nalidixic Acid (agar)
DNA: Deoxyribonucleic acid
EQUC: Expanded Quantitative Urine Culturing
GARD: *Gardnerella vaginalis* selective agar
HCCA: α-Cyano-4-hydroxycinnamic acid MAC MacConkey (agar)
MALDI-TOF: Matrix-assisted laser desorption ionization time-of-flight mass spectrometer
MRS: De Man-Rogosa-Sharpe (agar)
PCR: Polymerase chain reaction rRNA Ribosomal ribonucleic acid
RUTI: Recurrent urinary tract infections
TSA: Tryptic Soy (agar)
TSB: Tryptic Soy (broth)
UTI: Urinary tract infections
UTIC: CondaChrome® UTIC (agar)
UTIr: Urinary tract infections in postmenopausal women revisited (project)

## Declarations

### Authors’ contributions

Conceptualisation: FDE, PGM, MGJdV

Methodology: FDE, MDH, CvdZ, TV, GW, MJC, KLB, JKB, PGM, TH

Supervision: RSE, MGJdV

Visualisation: FDE, MGJdV

Writing – original draft: FDE, MGJdV

Writing – review & editing: RSE, MFS, AJW, JHHM, MGJdV

