## Supplementary Information for "An expanded urine culturing workflow to cultivate and characterize diverse urobiome isolates"

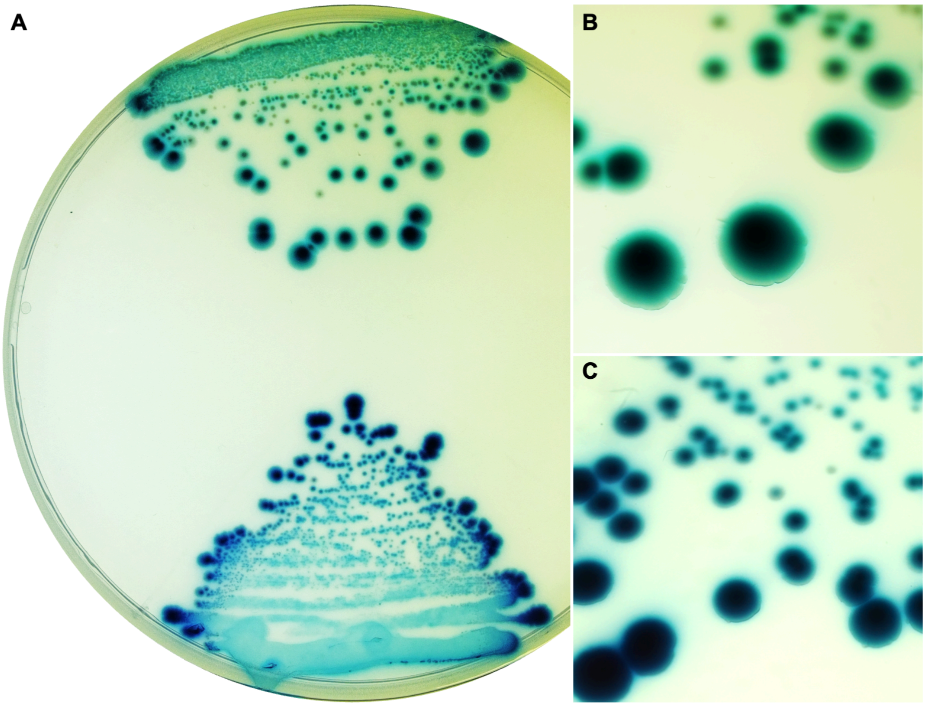


**Figure S1.** Morphologies of *E. faecalis* and *E. faecium* isolates grown on chromogenic agar (UTIC), following incubation for 3-4 days at 37ºC under 5% $CO_2$ conditions. **A** *E. faecalis* (top) and *E. faecium* (bottom) streaked on agar as viewed with a naked eye. **B** Macro image of *E. faecalis* colonies. **C** Macro image of *E. faecium* colonies.

**Table S1.** Morphological descriptions of Gram-positive urobiome taxa not previously described on Chromogenic (UTIC) agar, and not pictured in Figure 1 following incubation for 3-4 days at 37ºC under 5% $CO_2$ conditions. Taxa which have not yet been morphologically described may have similar appearances to those listed here. Species from the genera *Corynebacterium* and *Staphylococcus* are not pictured in Figure 1 due to high morphological variability and lack of species-level replicates. *Winkia* is not pictured in Figure 1 due to its very small colony size and difficulty of its color to show up on camera.

| Taxa | Colony Color | Colony Morphology | Agar Origin |
|---|---|---|---|
| *Corynebacterium* spp.<br>*C. freneyi*<br>*C. hesseae*<br>*C. pyruviciproducens*<br>*C. tuberculostearicum* | Cream | Variable size and shape, scanty | CNA |
| *Staphylococcus* spp.<br>*S. capitis*<br>*S. epidermidis*<br>*S. haemolyticus*<br>*S. hominis*<br>*S. lugdunensis*<br>*S. pasteuri*<br>*S. pettenkoferi*<br>*S. saprophyticus*<br>*S. simulans*<br>*S. ureilyticus* | Cream | Medium-large, spread out, variable shades of cream ranging from yellowish to pinkish | CNA |
| *Winkia* spp.<br>*W. neuii*<br>*W. anitrata* | Magenta | Punctiform, circular, opaque but low color intensity | CNA, TSA |

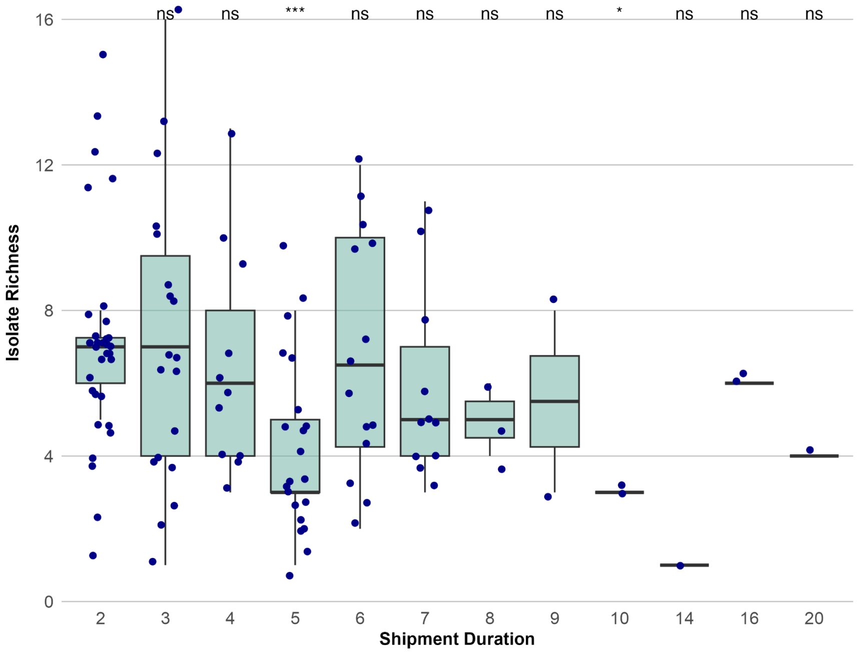


**Figure S2. Isolate richness at different shipment durations across urine samples, including all samples.** Isolate richness is defined as the number of morphologically distinct isolates as described on UTIC agar obtained from one urine sample (*n* = 108 urine samples). Shipment duration is defined as the number of days between urine sample collection and starting the culturing workflow. (ns = not significant, $*p < 0.05$, $**p < 0.01$, $***p < 0.001$).
